# Rubber needs to be included in deforestation-free commodity legislation

**DOI:** 10.1101/2022.10.14.510134

**Authors:** Eleanor Warren-Thomas, Antje Ahrends, Yunxia Wang, Maria M H Wang, Julia P G Jones

## Abstract

Natural rubber production uses increasing amounts of land in the tropics and is linked to deforestation. There is debate as to whether current legislative proposals to reduce the import of deforestation-linked commodities into the EU, US and the UK will include rubber. Globally, sustained growth in demand is chiefly driven by tyre production, linked to rising freight and passenger transport flows. Yields of natural rubber remain static, meaning increased plantation area will be required: 2.7 – 5.3 million ha of additional harvested area could be needed by 2030 to meet demand. In order to prevent further deforestation and associated biodiversity loss, millions of smallholder growers producing the majority of rubber globally need support to increase production from existing plantations and close yield gaps, without undermining long-term sustainability through soil or water degradation. Rubber should also be included in legislative proposals to reduce deforestation in supply chains to avoid undermining the impact of these ambitious initiatives on forest loss globally.

## Main text

Avoiding tropical deforestation is critical to protect biodiversity, address climate change, protect ecosystem service delivery, and support indigenous peoples (Lyons-White et al., 2020). As conversion to agricultural land is a key driver of forest loss, there are increasing initiatives to eliminate deforestation from the supply chains of agricultural commodities (Lyons-White et al., 2020; Seymour & Harris, 2019). Legislative proposals are currently under consideration in the EU (Directorate-General for Environment, 2021b), the US (S.2950 - 117th Congress (2021-2022): FOREST Act of 2021, 2021) and the UK (DEFRA, 2021) to regulate the import of deforestation-linked commodities.

Natural rubber *(Hevea brasiliensis)* is essential for the manufacture of vehicle and airplane tyres (comprising 70% of global natural rubber consumption (Laroche et al., 2022)), medical equipment, prophylactics and sportswear (Warren-Thomas et al., 2015). Some tyre companies have made zero-deforestation commitments, and a voluntary sustainability initiative, the Global Platform for Sustainable Natural Rubber (GPSNR) works to address deforestation alongside other sustainability concerns (https://sustainablenaturalrubber.org). However, only a small fraction of the rubber industry has voluntarily committed to avoiding deforestation.

The EU proposal for a regulation on deforestation-free products will require businesses placing goods containing specific named commodities on the EU market to show that products were not produced on land deforested or degraded after 31 December 2020 (Directorate-General for Environment, 2021b). Six commodities are currently included in the proposal (soy, leather, beef, oil palm, cocoa, coffee; (Directorate-General for Environment, 2021a)), but while rubber was initially assessed for inclusion, it was omitted based on an impact assessment (Directorate-General for Environment, 2021c). This assessment re-analysed published estimates for embodied deforestation risk in EU supply chains (Pendrill et al., 2019), but the authors of the original study found this to be flawed, and concluded that there is no empirical basis to exclude rubber (Persson et al., 2021). Similar legislative proposals under earlier stages of consideration in the US and the UK may or may not include rubber. While the EU proposal is a very positive step, despite the known challenges of meeting sustainability standards in smallholder-dominated supply chains (Grabs et al., 2021; Lyons-White et al., 2020), omission of rubber, even if only in the near term, could risk undermining the efficacy of these legislative efforts to avoid deforestation.

Here, we provide evidence that demand for rubber in the coming decade will likely lead to a further expansion of plantation area, and that recent area expansion has been linked to deforestation in the tropics. We argue that rubber should be included in the proposed EU regulation alongside other forest-risk commodities, and that more support is needed for smallholder farmers already cultivating rubber to sustainably improve their yields, to support livelihoods without increasing the global footprint of rubber plantations.

### Increasing rubber demand was met by expansion of plantation area 2010-2020

Global rubber production has increased steadily over the past decade, but as yields per hectare have been mostly stable, this has been achieved through an increase in plantation area (FAO, 2022) (**Figure 1**, **Figure S1, Supplementary Methods**). A series of feedback loops among drivers of demand, supply, price, area expansion, crude oil price, and supply constraints together explain why rubber area continues to increase despite relatively low prices, and reported problems for smallholder profitability in some locations (Ali et al. 2021).

**Figure 1:**
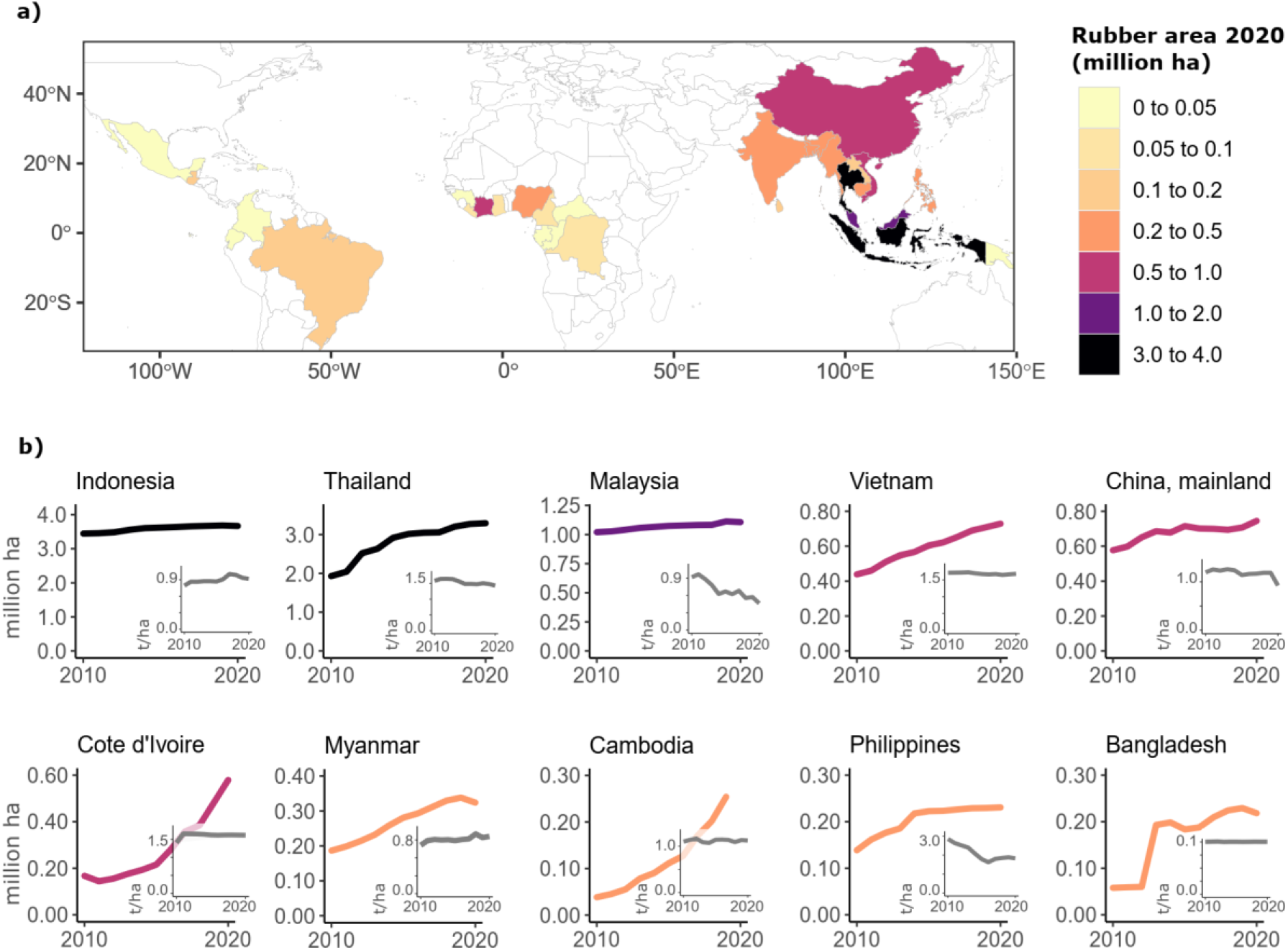
a) Harvested area of natural rubber per country in 2020. b) Harvested area of natural rubber from 2010 to 2020 for the top ten countries for absolute rubber area increase over the same period (colour of the main graph line reflects the 2020 total shown on the map, and the key per country; subplots per country show yields in metric tons per hectare). All data from FAOSTAT, except for government estimates reported for Lao PDR (Supplementary Methods).

Global harvested area increased by 3.3 million ha between 2010 and 2020, bringing the total to 12.8 million ha (FAO, 2022). This falls at the higher end of expansion projections based on demand increase between 2010 and 2018, under multiple assumptions of intensification, yields, and displacement of rubber area with oil palm (0.9 – 4.2 million ha) (Warren-Thomas et al., 2015).

Most rubber-producing countries increased their harvested rubber area between 2010 and 2020 (FAO, 2022) (**Table S1**). The largest increases were in Thailand (1.4 million ha) and Cote D’Ivoire (0.41 million ha), followed by more than 0.2 million ha in Vietnam, Cambodia, and Indonesia, respectively. Only one country reduced its rubber area (India). These data represent mature plantations, not total planted area: as rubber takes 5-10 years to mature, this expansion only represents areas planted up to 2015.

### Rubber expansion has been linked to deforestation

Globally, 80% of rubber is grown by smallholders (Laroche et al., 2022). Rubber plantations have already been linked to deforestation in Southeast Asia, involving both smallholders and companies operating larger industrial-scale estates (Warren-Thomas et al., 2015), with impacts on biodiversity, carbon, and people **(Supplementary Text)**. However, patterns of smallholder-driven land cover and land use change to rubber are more complex than change for some other plantation crops, such as oil palm, which tend to be established in larger contiguous blocks. This partly explains why detection of rubber plantations from earth observation satellites is more challenging than for other tree plantations (Ye et al., 2018), and no analysis has yet quantified deforestation for rubber at the global scale. However, multiple sources of evidence show that a substantial proportion of recent rubber area expansion has involved deforestation, particularly in Asia.

A recent regional-scale analysis of mainland Southeast Asia quantified forest conversion to rubber up to 2014 using remote-sensing (Hurni & Fox, 2018). Data provided by the authors indicates that of 2.8 million ha of rubber area established between 2003 and 2014, 1.8 million ha replaced forest: 49% of this was in Cambodia, 18% in Vietnam, 15% in Laos, 8% in China, 6% in Vietnam, and 4% in Thailand. The remainder was established on areas classed mostly as annual cereal crops, but also grass and shrubs. Another regional assessment for seven countries in Asia, Africa and South America overlaid rubber plantation outlines with tree loss and gain maps, and detected 2.1 million ha of deforestation between 2001 and 2015 (Dow Goldman et al., 2020).

Local-scale case studies also describe land use and land cover change to rubber in the past decade (44 published reports, **Figure 2, Data S1**). In some regions rubber is replacing swidden systems (shifting cultivation), which include a mosaic of secondary forest patches that represent ‘invisible’ reservoirs of biodiversity (Padoch & Pinedo-Vasquez, 2010). Where swidden agriculture for rice is replaced with rubber, as has occurred in many parts of Southeast Asia, there is also a risk of food crop displacement to forest frontiers, and impacts on food security. This has been shown in Laos, where replacement of swidden with rubber displaced demand for food crops, leading to clearance of intact forest (Hurni & Fox, 2018). In some parts of mainland Southeast Asia, insular Southeast Asia and West Africa, old growth forests have been converted (**Figure 3;** in Cambodia, 20,000 - 100,000 ha of forest were converted to rubber each year 2010 – 2015; (Grogan et al., 2019); see also case studies in **Data S1**), with well documented negative impacts on biodiversity (Gibson et al., 2011; Prabowo et al., 2016). The introduction of agro-industrial rubber plantations into areas with formerly low human population density also introduces a risk of associated deforestation through increased food and fuel demand for plantation workers and their families (**Supplementary Text**).

**Figure 2:**
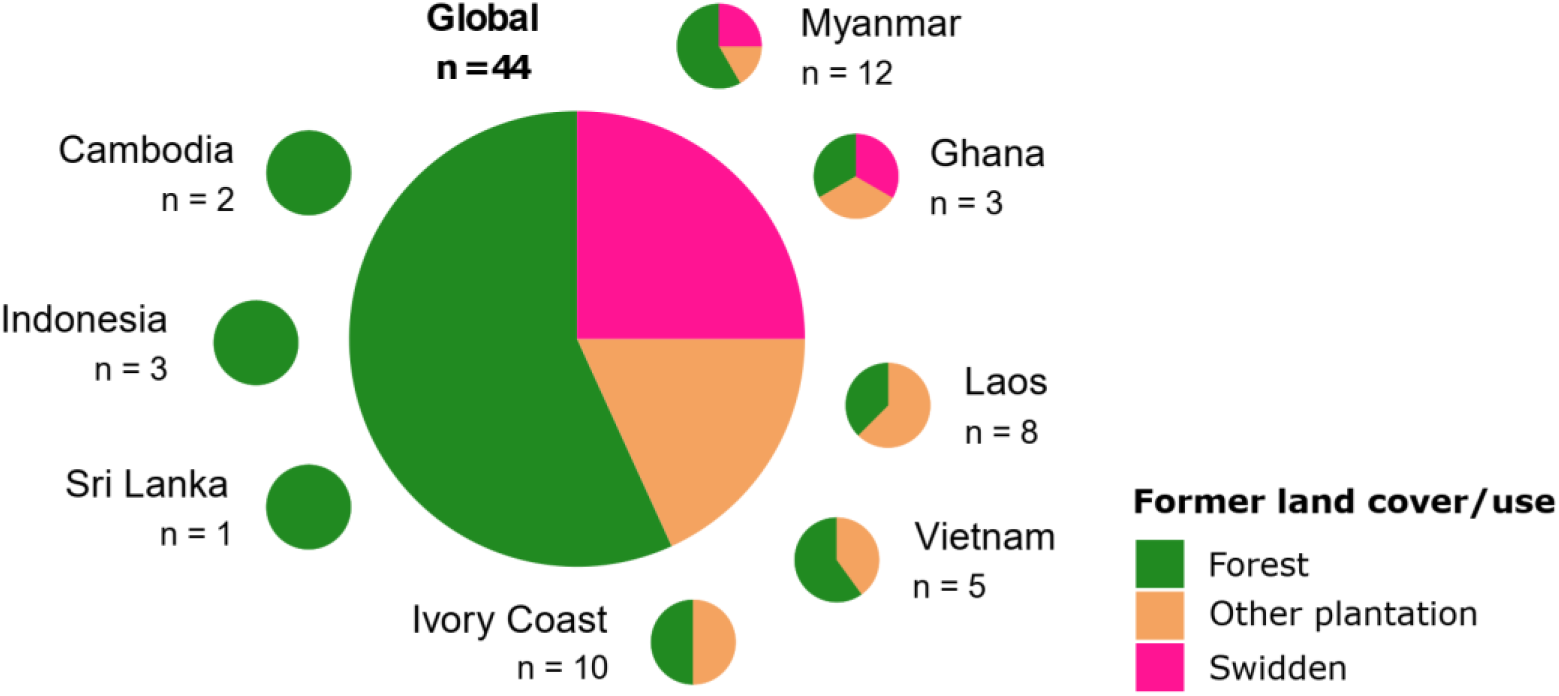
Number of published case studies (2009 – 2022) reporting conversion of forest, swidden or other plantation land to rubber plantations in the past decade (case study detail provided in **Data S1**).

**Figure 3.**
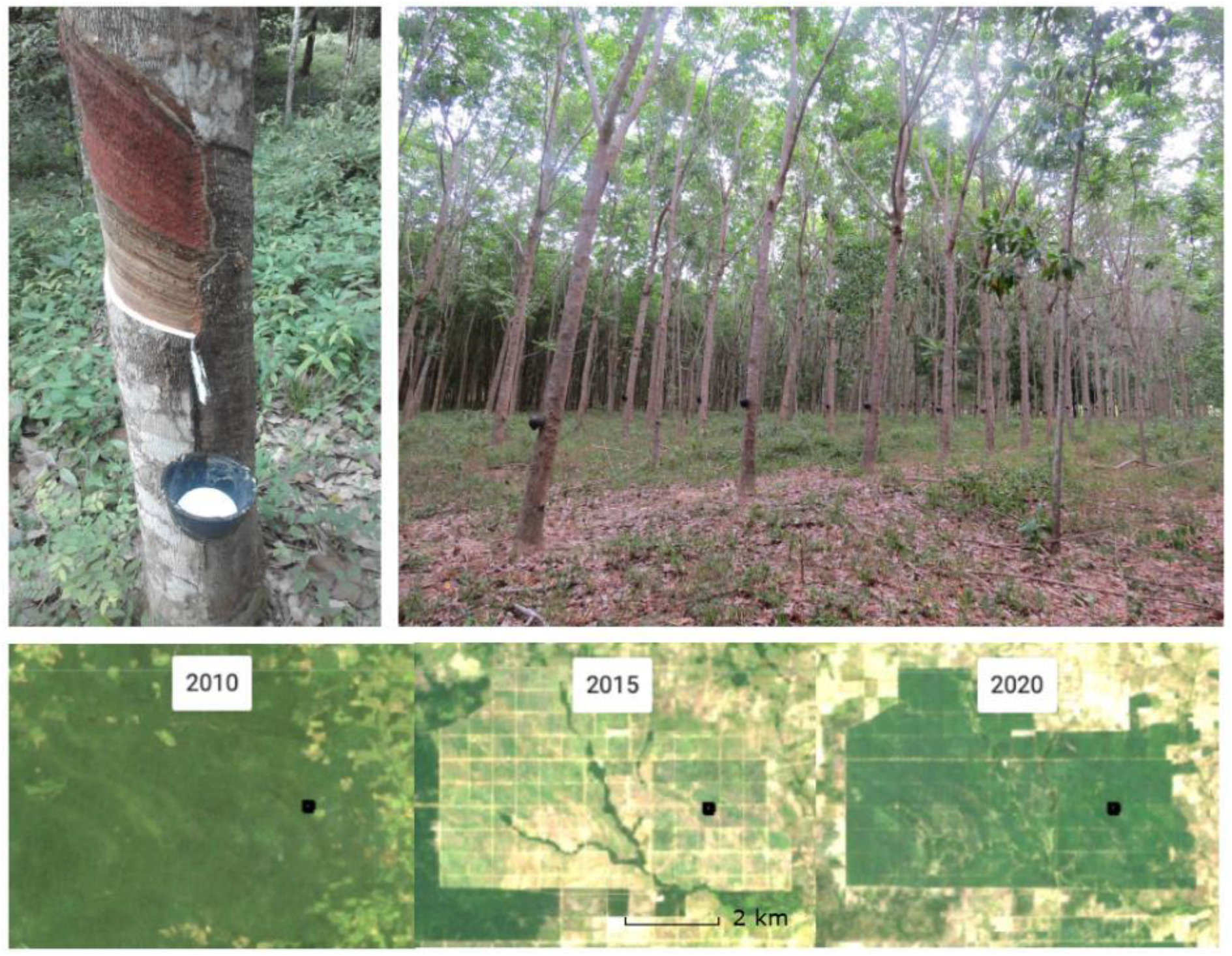
Top left: tapping a rubber tree in Southern Thailand. Top right: smallholder rubber plantation in Thailand (photograph credits: Eleanor Warren-Thomas). Bottom: Landsat true color annual composite images (courtesy of the U.S. Geological Survey) of intact forest (in 2010, R/G/B = Landsat-5 Band 3/2/1), forest clearance (by 2015, R/G/B = Landsat-8 Band 4/3/2) and rubber plantations (by 2020, R/G/B = Landsat-8 Band 4/3/2) in Northern Cambodia (Oddar Meanchey Province).

**Figure 4.**
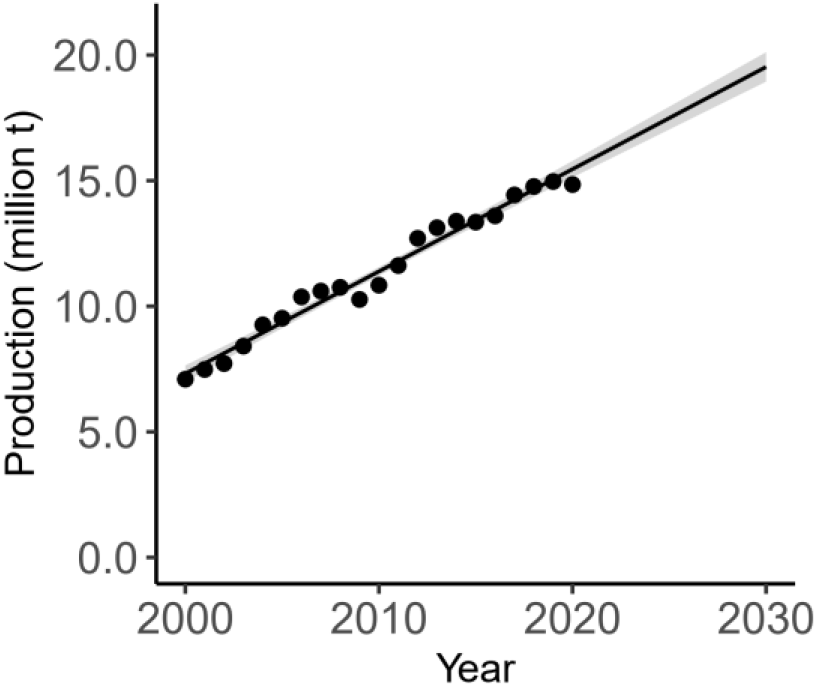
Historical rubber production globally 2000-2020, with fitted linear model of production in response to year, and model prediction to 2030. Data from FAOSTAT (model structure, code and full references in Supplementary Materials)

In West Africa, there is also a complex interaction with deforestation for cocoa, as rubber replaces old cocoa, and cocoa is expanded into the forest frontier. Land cover and land use change for rubber in West and Central Africa warrants particular attention, given that many nations still have high forest cover (Lyons-White et al., 2020), and the EU increasingly seeks to source rubber from this region ((Laroche et al., 2022) and **Supplementary Methods**).

Finally, according to government statistics, forest cover loss and rubber area gain have often coincided at the subnational level in the top two rubber-producing countries, indicating locations where rubber may be associated with increased pressures on forest (**Supplementary Methods**). In Indonesia, between 2014 and 2022, 10 of the 17 provinces where rubber area increased also had a loss of forest area; this was concentrated on the islands of Sumatra and Borneo, where forest loss was reported in five of five, and three of four provinces with rubber gains, respectively (**Data S2**). In Thailand, between 2012 and 2018, rubber area increased in 46 provinces, of which 23 had forest loss; this was particularly concentrated in the northeastern region, where all 20 provinces reported rubber gains (**Data S3**).

### Rubber demand will increase to 2030

Globally, demand for natural rubber is likely to continue rising. Recent contractions in demand due to a global semiconductor shortage and reduced vehicle sales during the COVID-19 pandemic are likely to be of a short-term nature (World Bank Group, 2021). Global freight transport and passenger flows are predicted to triple by 2050 (International Transport Forum, 2019). Interestingly, sustainability concerns mean there are also industry efforts to increase the proportion of natural rubber used in tire manufacturing: petroleum-based synthetic compounds are currently used in combination with natural rubber (for example, passenger vehicle tires are <50% natural), but there are now efforts to reduce reliance on synthetics (Laroche et al., 2022). Historical trends of expansion, and static yields per unit area, mean increased demand is likely to be met with expansion: so how much additional area could be needed to meet demand by 2030?

Estimates of future demand can be made using projections from rubber industry analysts (**Table S2**), which indicate natural rubber demand would reach 17.2 million metric tons by 2030 (an increase of 4.6 million metric tons from 2020). An alternative approach is to project future production by extrapolating historical trends over time using a simple linear model, based on observed data 2000-2020 (FAO, 2022). Production in 2030 is then predicted to be 19.5 million metric tons (**Table S3**). The additional harvested rubber area required to meet the projected 2030 demand can be estimated using average yields for 2010 to 2020 from the highest yielding (Vietnam, 1.69 metric tons/ha) of the top ten producing countries (by increase in absolute area 2010-2010), or the lowest yielding (Indonesia, 0.89 metric tons/ha). This shows that meeting projected 2030 demand could require 2.7 – 5.3 million ha of additional harvested rubber area relative to 2020 (**Table S2, Table S3**). This expansion is likely to come into direct conflict with priority areas for biodiversity conservation (Wang et al., 2020).

Disease and climate change risk may further add to land demand for rubber by reducing productivity of existing plantations: pathogens are an old problem in rubber plantations, but *Pestalotiopsis* is currently reducing yields by 25% across ~0.4 million ha of rubber in Indonesia, and multiple leaf-fall diseases are affecting China, Indonesia, Thailand, Malaysia, Sri Lanka and India (Pinizzotto et al., 2021). Changing weather patterns are already impacting plantations across South and Southeast Asia: prolonged dry seasons can reduce the risk of fungal disease but reduce rubber tree growth and latex flow, warmer winters in China are linked to increased fungal damage, while intensified rainfall reduces available tapping days and causes flooding that prevents harvest **(Supplementary Text)**. Moreover, many existing plantations have been established in sub-optimal locations prone to storm and erosion risk, which are likely to become increasingly marginal as climate change progresses, further limiting production from existing rubber area (Ahrends et al., 2015).

### Sustainable intensification

Meeting natural rubber demand without incurring deforestation risk will require increased production on the already-vast areas planted with rubber, without undermining long-term sustainability through soil degradation or water pollution. Existing evidence shows constraining land availability for agricultural expansion by strengthening voluntary and regulatory zero-deforestation commitments can increase investment on existing cultivated land, and increase yields, by redirecting investment from expansion to intensification (Lyons-White et al., 2020). However, smallholders can be highly reliant on international supply chains but have limited capacity meet the demands of sustainability initiatives (due to financial, land tenure and/or knowledge constraints); safeguards and support are needed to ensure continued market access and avoid displacement by other actors more able to adapt (Grabs et al., 2021). Volatility in rubber prices (**Figure S2**) makes it difficult for smallholders to invest in increasing yields, while high labour costs or shortages can mean they switch to alternative crops: price guarantees may help (**Supplementary Text**). Rubber smallholders also need further technical support to increase yields within resilient production systems by adopting climate- and disease-resilient rubber clones, and best management practices (Pinizzotto et al., 2021). Further research and development of intercropping and agroforestry practices for rubber could also offer multiple benefits, including improved water and soil management (Wang et al., 2021), while also supporting some elements of biodiversity (Prabowo et al., 2016; Warren-Thomas et al., 2019).

High rubber prices following supply constraints in 2011 were directly associated with industrial-scale deforestation for rubber in Cambodia (**Figure S2; Figure 3,** (Grogan et al., 2019)). Meanwhile, particularly in the expansion frontiers of Northern Laos and Central and West Africa, rubber remains an attractive investment even when prices are relatively low. Deforestation regulations applied to rubber could help prevent rapid expansion onto forests in response to high prices in future, if sufficiently widespread (**Supplementary Text**).

### Inclusion of rubber in zero-deforestation legislation

Among the many complex issues raised by zero-deforestation commitments (including what constitutes forest, traceability and effective implementation; we highlight specific challenges for rubber in the **Supplementary Text**), leakage across space, among ecosystems, and between commodities is a particular challenge (Lyons-White et al., 2020). Land use change to rubber has documented interactions with other forest-risk commodities, including cocoa (West Africa), oil palm (insular and southern southeast Asia) and rice (northern mainland southeast Asia), as well as increasing the total demand for land in the tropics. Together this means that rubber’s exclusion from zero-deforestation legislation increases the risk of between-commodity leakage effects and would decrease overall efficacy of the regulations.

We recognize the potential risks for smallholder farmers posed by deforestation-free legislation, if support and safeguards are not provided. Unintended consequences may arise, such as the exclusion of smallholders and smaller businesses from markets and landscapes, and the placing of additional challenges onto weaker economies with insufficient capacity provide evidence for deforestation-free production (Grabs et al., 2021). Much more work needs to be done to realize effective and equitable zero-deforestation commitments, particularly to understand the impact on people dependent on agriculture in landscapes containing tropical forests (Lyons-White et al., 2020), and to ensure that smallholder farmers are supported to find alternative strategies to forest clearance (Seymour & Harris, 2019). However, the necessity of slowing deforestation is clear. Omitting rubber from the deforestation-free commodity legislation risks undermining action on forest loss globally, and should be reconsidered.

## Acknowledgments

We thank Chris West for useful critical comments on a draft of this paper.

## Author contributions

Conceptualization: EWT

Data curation: EWT

Formal analysis: EWT

Methodology: EWT

Investigation: EWT, JPGJ, MMHW, AA, YW

Visualization: EWT, AA, YW

Project administration: EWT

Writing – original draft: EWT

Writing – review & editing: EWT, JPGJ, MMHW, AA, YW

## Data and materials availability

Data S1 to S4 are available on request from lead author.

## Supporting Information

Supplementary Methods

Supplementary Text

Figures S1 to S2

Tables S1 to S3

Supplementary Code 1

Supplementary References

## Supporting Information for

### Other Supplementary Materials for this manuscript available on request from lead author

Data S1: Case studies of land use and land cover change for natural rubber 2010 – 2020

Data S2: Sub-national data on rubber and forest area per province in Indonesia

Data S3: Sub-national data on rubber and forest area per province in Thailand

Data S4: FAOSTAT data on rubber area, production and yield used as input for Supplementary Code 1 to project rubber production to 2030

## Supplementary Methods

### Rubber consumption by sector and by EU

Estimates of the amount of natural rubber that is used for the tire manufacturing industry have been made by the World Bank (“two-thirds”) (World Bank Group 2021) and the Rainforest Alliance (70%) (Millard 2019).

The EU (including the UK) accounted for 9% of global natural rubber consumption in 2017, sourced chiefly from Indonesia, Thailand and Cote D’Ivoire. In 2016, EU demand for tires was estimated to require 0.59 million ha of harvested rubber area (5% of global total; range 0.34 and 1.4 million ha due to uncertainty and opacity of trade flows) and formed a particularly high proportion of the total harvested area grown in Cambodia, Cote D’Ivoire, Guinea and Cameroon, reflecting a policy to increasingly source from the African continent (Laroche et al. 2022).

The European Tyre & Rubber Manufacturers’ Association (ETRMA) submitted evidence to the EU as part of the consultation process for the proposed regulation, including statistics from the International Rubber Study Group (IRSG) and Eurostat for the year 2017 (ETRMA 2019). On the EU’s role in natural rubber consumption in 2017: the EU (EU28 including the UK) was responsible for 9% of total global natural rubber consumption (China 37%, India 8%, US 7%, Japan 5%, Thailand 5%, rest of the world 29%). EU imports were sourced from Indonesia (32%), Thailand (19%), Cote D’Ivoire (19%), Malaysia (12%), Vietnam (8%), Cameroon (2%), and 8% from other countries. Imports from Cote D’Ivoire are reported to be increasing. The statistics for 2020 had changed slightly, with EU and UK combined consumption representing 8% of the global total, and China 43%; imports were sourced from Indonesia (28%, a decrease), Thailand (21%, a decrease) and Cote D’Ivoire (23%, an increase) (ETRMA 2021).

One study conducted detailed analysis of EU consumption of natural rubber for mobility (i.e. tires for bicycles, motor vehicles and airplanes) by conducting input-output analysis of trade flows to track tire demand to source countries for EU countries (including the United Kingdom) (Laroche et al. 2022). A matrix of EU natural rubber consumption (by weight, for mobility) by source country showed that 26% of consumption was sourced from Indonesia and Thailand respectively, 14% from Cote D’Ivoire, 6% from Malaysia, 11% from China and 7% from Vietnam. This rubber demand for EU mobility then translated into a land footprint of 0.594 million ha total (5% of global area, but uncertainty in some values and opacity of trade flows mean this value could be as low as 0.342 million ha, and as high as 1.6 million ha. The land footprint (area of harvested rubber) was distributed among Indonesia (32%), Thailand (23%), Malaysia (11%), China (11%) and Cote D’Ivoire (10%). As a proportion of the total harvested rubber area consumed per producer country, EU consumption accounted for 25% of the total harvested area in Cambodia, and >15% in Cote D’Ivoire, Guinea and Cameroon.

### Data on rubber harvested area, production and yields

All data on rubber harvested area and production reported in the main text were downloaded from FAO production statistics available via FAOSTAT (FAO 2022) in January 2022, using the item “Rubber, natural”. We note the potential for data inaccuracies related to differences in methods and quality of national reporting among countries of both rubber production and forest loss statistics to the FAO. However, whilst the figures and their national disaggregation may be imprecise, the overall trends are likely to be captured in these data given that the figures are corroborated by various other national and regional studies and reports.

### Projections of future rubber demand

Industry expert estimates of future rubber demand are published quarterly and annually by the IRSG (IRSG 2022) and have been used in previous studies of rubber expansion (Wang, Carrasco, and Edwards 2020; Warren-Thomas, Dolman, and Edwards 2015). The projections for 2021 onwards in this study were taken from reporting of the IRSG’s headline findings by Tires & Parts News published on January 19, 2022 (Tires & Parts News 2022). The IRSG predicted that after contraction from 2019-2020, total demand for both synthetic and natural rubber 2020-2021 was expected to increase by 9.4% to reach 29.57 million t (metric tons) in 2021, of which natural rubber comprised 47%. Total rubber demand was then predicted to increase by 3.6% 2021-2022, and subsequently by an average 2.3% year-on-year from 2023-2030. Assuming the proportion of total demand met by natural versus synthetic rubber remains the same as in 2020, natural rubber demand to 2030 can be projected (**Table S2**).

Historical trends in rubber production (2000 – 2020) were taken from FAOSTAT, and fitted against year in a simple linear model (**Table S3**) using R (**Supplementary Code 1**), following the method used by a previous study (Parra-Paitan and Verburg 2022).

### Subnational rubber and forest area changes in Thailand and Indonesia

Subnational data on rubber plantation and forest area change for Thailand and Indonesia were taken from government statistics. In Thailand, forest cover statistics for each province are published by the Department of National Parks, Wildlife and Plant Conservation (Department of National Park, Wildlife and Plant Conservation (DNP) 2022); data published in reports covering the years 2012-2013 and 2018-2019 were used for analysis. Rubber plantation area data are published in annual tables by the Office of Agricultural Economics (Office of Agricultural Economics 2022); data for the years 2012 (the earliest available) and 2018 were used for analysis. In Indonesia, rubber plantation area data are published by the Badan Pusat Statistik (Central Agency of Statistics) (Badan Pusat Statistik 2022) and forest cover data tables per province are published by the Ministry of Environment in an annual report (Ministry of Environment and Forestry 2021), for which data on all forest types (permanent and conversion, secondary and primary) were included. Rubber area data for two provinces (Kalimantan Timur, Kalimantan Utara) were combined (under Kalimantan Timur) to match forest data.

## Supplementary Text

### Old-growth forest, secondary forest and swidden conversion to rubber: carbon, biodiversity, social and economic outcomes

It is widely understood that intact forest ecosystems are essential for biodiversity conservation in tropical regions, as many species are completely dependent on them, but secondary and logged forests also support much greater biodiversity value than monocultural plantations, including rubber (Barlow et al. 2007; Gibson et al. 2011; Prabowo et al. 2016). Simulations of rubber expansion to 2027, based on industry projections of future demand, found that if expansion takes place in the most climatically suitable locations for rubber, 74 forest-dependent species could lose >= 10% of their range, and six species could lose >=50% of their range due to forest loss (Wang et al. 2020). This does not account for interactions with other crops vying for space in the same bioclimatic zones, such as oil palm (Warren-Thomas et al. 2015). Natural forest conversion to rubber also results in net carbon emissions (Warren-Thomas et al. 2018).

Rubber expansion is still considered a major driver of deforestation in landscapes important for biodiversity conservation in Southeast Asia, second only to rice: an expert opinion survey of conservation managers looking after landscapes important for biodiversity globally (working for the Wildlife Conservation Society, and managing not just protected areas but surrounding landscapes), found that rubber was perceived as a driver of forest loss in six landscapes in Cambodia, Malaysia, Myanmar, Thailand and India (Jayathilake et al. 2021). Given historical patterns of widespread rice replacement with rubber in many countries in Asia (e.g., Thailand), these findings further highlight the ongoing risk of food crop displacement to forest frontiers.

Where agro-industrial plantations have been developed in formerly forested landscapes with low human population densities, a rapid increased in food and fuel demand to support plantation workers and their families can impact surrounding landscapes beyond the boundaries of plantations. This has been reported in eastern Cambodia (Fox et al. 2018) and raised as a concern for the development of plantations in Central African countries (Cameroon, Gabon, Republic of Congo and Democratic Republic of Congo), based on experiences with mining operations (Feintrenie 2014).

Swidden, or shifting agriculture, landscapes comprising mosaics of farmed areas, grazing, fallows of variable age and secondary forest patches, represent ‘invisible’ reservoirs of biodiversity that are often ignored in debates about biodiversity conservation (Padoch and Pinedo-Vasquez 2010). Forest patches in swidden systems likely play a key role in supporting biodiversity at the landscape scale, relative to conversion to rubber monoculture. The impact of conversion of swidden to rubber on carbon emissions are complex and context dependent (Fox, Castella, and Ziegler 2013).

From a social and economic perspective, switching from subsistence agriculture to cultivating rubber can bring huge economic benefits for farmers, but the transition to rubber plantations has also had social repercussions in some places. Land tenure, exploitative contracts, and food security issues are reported from parts of Asia and West Africa. Land tenure problems and exploitative contracts with growers have been reported, for example in northern Vietnam (Dao 2018; van Vliet et al. 2012), southern Laos (Kenney-Lazar 2012), Cambodia (Fox and Castella 2013), and Myanmar (Kenney-Lazar et al. 2018). There are also concerns about rubber replacing food production, risking food security in Cote D’Ivoire and Ghana (Akmel 2018; Owusu and Ruf 2015), as well as concerns from non-governmental organizations about labor rights in some Asian and African countries (Aidenvironment 2020).

Agroforestry practices may improve biodiversity value of rubber plantations, whether complex ‘jungle’ systems that have very low rubber yields, or simple high-yielding systems that include additional crops or have high levels of natural vegetation understory (Clough et al. 2016; Prabowo et al. 2016; Warren-Thomas et al. 2020). Agroforestry, or diversified systems, can also provide multiple benefits for soils and water management on rubber farms, alongside benefits for farmers (Wang, Warren-Thomas, and Wanger 2021).

### Disease and climate change risk in existing rubber plantations

Modelling of climate change impacts on rubber suitability in mainland Southeast Asia has indicated an expanding area of suitability through climate warming and reduced risk of frosts (Golbon, Cotter, and Sauerborn 2018). However, other climate risks to plantations (storm damage from typhoons, soil erosion from intensified rainfall events, and water stress from drought) are predicted to decrease the area of suitability in the same region (Ahrends et al. 2015). Disease risk response to climate change has not yet been modelled.

Rubber plantation managers across Asia (Indonesia, Thailand, Malaysia, Sri Lanka, India) are reporting increasing damage from fungal diseases, and some plantation managers are using fungicidal fog treatments to reduce damage, with increased fungal infection linked to wetter rainy seasons (Pinizzotto et al. 2021). Sulphur dust treatments are also commonly used; both treatments cause environmental pollution(Liyanage et al. 2016). Work in China indicates that warmer winters are associated with increased prevalence of powdery mildew disease due to changes in leaf phenology (rubber trees drop their leaves during the winter/dry season) (Zhai and Xu 2022). Meanwhile, a new fungal pathogen of rubber trees has recently been identified in Thailand (Pornsuriya et al. 2020). Fungal pathogens can reduce rubber yields by 12 – 45% (Liyanage et al. 2016; Pinizzotto et al. 2021).

### Sustainably increasing production on existing rubber farms/understanding continued expansion

Relatively low prices currently mean that despite net expansion of rubber area, some smallholders are giving up cultivation, or struggling to afford necessary inputs to maintain yields. Farmer responses to price fluctuations vary by location. In southern Myanmar, farmers reduce rubber production but retain trees when prices are low (Vagneron et al. 2017). In contrast, smallholders in Malaysia have felled rubber trees for timber value and switched to other crops (such as oil palm) or off-farm activities (Ali et al. 2021).

In Cambodia (Grogan et al. 2019) and Vietnam (Kissinger 2020) expansion of rubber area is clearly linked to international prices, with harvested area of rubber in Vietnam contracting following price crashes in 2016, while Cambodian industrial plantations are required by law to keep planting, but incorporate cashew and pepper to boost income (Hurni and Fox 2018).

In northern Laos, expansion of rubber showed characteristics of path-dependency or lock-in of land use transitions beyond an initial trigger (such as temporarily high prices (Junquera et al. 2020)). Here, differences in income between subsistence swidden cultivation and rubber are large, even when rubber prices are low, and people perceived prices as likely to increase again in future. Rubber was also perceived as a means to secure land tenure. In another main frontier of rubber area increases, Cote D’Ivoire, rubber smallholders report steadier (monthly) and greater income from rubber than from alternative plantation crops (coffee, cocoa, oil palm) following price declines for coffee and cocoa (Akmel 2018).

The GPSNR is working towards improving many issues around smallholder profitability, yields and vulnerability to price fluctuations through its capacity building workstream, which is focused on reducing yield gaps due to poor management practices (Maria Wang personal observations after attending GPSNR workshops). They are developing generalized “Good Agricultural Practices” to be shared with smallholders through a mobile phone app, some of which are based on management practices in Cote D’Ivoire (including reduced tapping intensity) which are contributing to relatively high yields compared to other countries. Other GPSNR strategies include providing training, high-yielding planting material and disease management resources in Indonesia, and expanding agroforestry practices in Thailand.

### Challenges for implementing zero-deforestation supply chains in the context of rubber

#### Forest definitions and indirect land use change (iLUC)

Forest definitions are a major sticking point for successful implementation of zero-deforestation: definitions vary among international organizations, nations and even landscapes, while in high forest-cover countries, there may be few options for agricultural development without forest clearance (Lyons-White et al. 2020). Logged, degraded and regenerating secondary forests support biodiversity and rapidly increasing carbon stocks, and form the majority of natural forest cover in some major rubber-producing landscapes, especially Vietnam (Phuc and Nghi 2014). Meanwhile, the definition of rubber plantations as planted forests or agriculture varies among countries: the FAO Global Forest Resources Assessment includes rubber trees as (planted) forest cover, but some countries count rubber plantations as an agricultural crop, and not all rubber-producing countries reported their rubber plantation area (FAO 2020).

In Sumatra, clearance of logged forest for rubber was permissible within zero-deforestation commitments made by Michelin, as forests were not classed as high conservation value or high carbon stock, despite supporting large mammals such as Sumatran elephants, and being adjacent to a national park (Otten et al. 2020). In Vietnam, most natural forest is secondary following widespread deforestation during the 20th century: these naturally regenerating secondary forests have considerable value for biodiversity (Meyfroidt and Lambin 2008) and rapidly increasing carbon stocks, but can legally be converted to rubber (Phuc and Nghi 2014). To have maximum benefit, and to be coherent with ambitious global forest restoration targets (secondary and logged forests are on trajectories of recovery to fully functioning forest ecosystems), definitions of deforestation may need to be broad, while forest cover loss within shifting agriculture (swidden) systems also needs careful consideration.

The role of rubber in indirect land use change, or leakage effects of eliminating deforestation from the rubber supply chain, also need to be carefully considered when assessing deforestation risk. Replacement of former cocoa plantations in Ivory Coast with rubber is linked to deforestation for new cocoa plantations (Ruf 2015), while forest clearance for subsistence farming has been detected following swidden conversion to rubber (Hurni and Fox 2018).

#### Traceability and transparency

For companies to implement zero-deforestation, whether voluntarily or to meet the demands of regulation, they need to be able to trace their source materials through complex supply chains. A key challenge is that buyers often import partly or fully processed products (mostly from China), rather than direct from source countries. For example, it is impossible to directly source rubber from Vietnam as a foreign company (Kissinger, Brockhaus, and Bush 2021). The role of China as the majority importer of natural rubber from southeast Asian growers is critical (China imports 40% of Malaysia’s rubber exports (Ali et al. 2021), and 40% of Vietnam’s (Phuc and Nghi 2014). Work in Sri Lanka, where 88% of domestic production is processed in-country, shows there is near-complete opacity between farm gate and processors, making it impossible for importers to know if they are linked to deforestation (Cho et al. 2022). These problems are magnified by the complexity of cross-border trade. Traceability tools specifically for rubber, such as RubberWay (https://rubberway.tech) have been developed to support voluntary private-sector supply chain tracing, while traceability and transparency is a major workstream for the Global Platform for Sustainable Natural Rubber, which has commissioned two reports identifying solution for the rubber sector (de Bonafos 2020; Cupit et al. 2020). Regulation could make such efforts mainstream, as for FLEGT (Directorate-General for Environment 2021).

**Figure S1.**
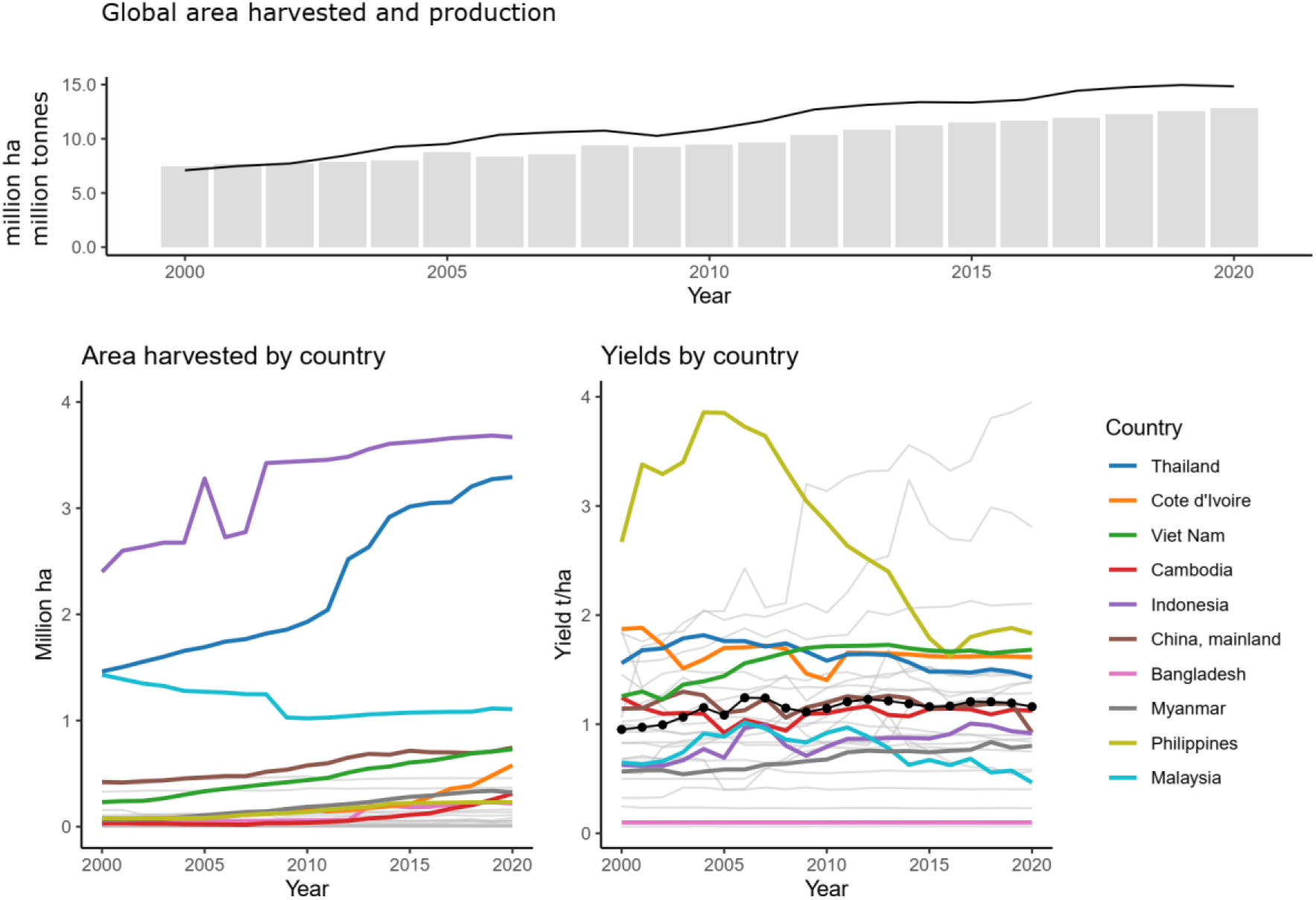
Global and national rubber area harvested (million hectares, bars), rubber production (million metric tons, line), area harvested per country (highlighted in color for the top 10 countries based on absolute area increase 2010-2020; million hectares; lines), and yields 2000 - 2020 (metrics tons per hectare; lines). While production and area have steadily increased since 2000, yields have remained relatively static (yields are calculated from harvested area and production by FAOSTAT). Yields above two metric tons per hectare are likely the result of incomplete reporting of harvested rubber area or production statistics (the two countries showing rapid increases from 2010 onwards are Guatemala and Mexico, which also produce latex from alternative species, so may not solely represent *Hevea brasiliensis*), while yields for Bangladesh of 0.1 metric tons per hectare are likely too low, and may represent under-reporting of production, or over-reporting of harvested area. Data from FAOSTAT (FAO 2022).

**Figure S2.**
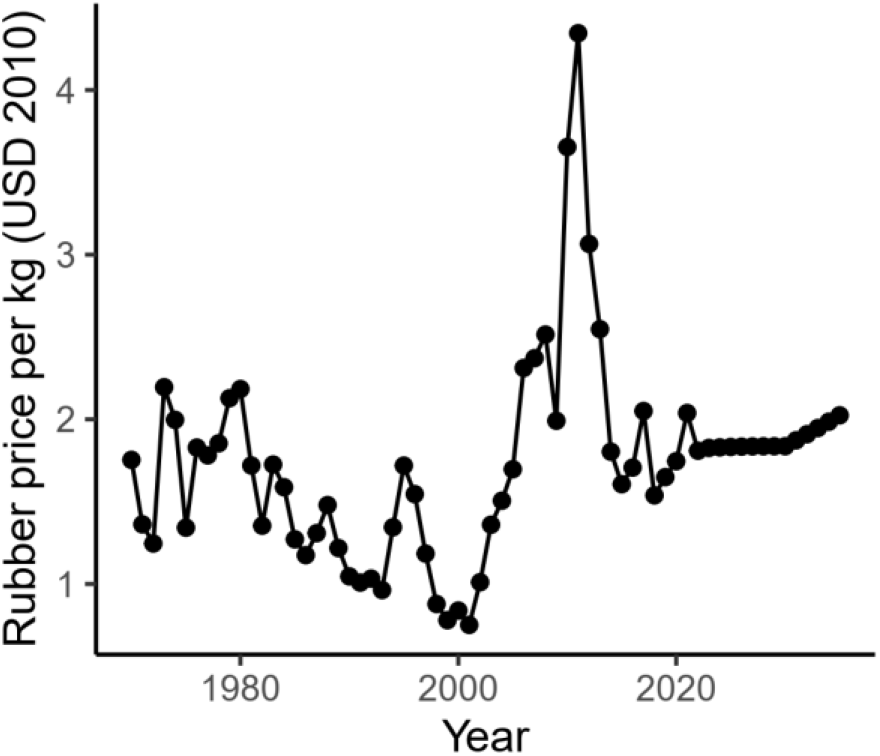
Price data in standardized 2010 USD from the World Bank Commodity Outlook (World Bank Group 2021) (historical to 2020, projected to 2020-2030). The peak price in 2011 stimulated rubber plantation expansion, but declining Chinese import volumes, production increases and falling crude oil prices meant prices subsequently declined (Vagneron et al. 2017).

**Table S1.** Top ten countries for increase in absolute rubber harvested area 2010-2020 reported to FAOSTAT (FAO 2022). Note: Lao PDR does not report data to the FAO, but government data indicates planted rubber area increased from 0.14 million ha in 2008, to 0.26 million ha in 2018, of which 0.12 million ha was being harvested (Smith et al. 2020).

| Country | Absolute change in harvested area 2010-2020 (million ha) |
| --- | --- |
| Thailand | 1.36 |
| Cote d'Ivoire | 0.41 |
| Viet Nam | 0.29 |
| Cambodia | 0.27 |
| Indonesia | 0.22 |
| China, mainland | 0.17 |
| Bangladesh | 0.16 |
| Myanmar | 0.14 |
| Philippines | 0.092 |
| Malaysia | 0.086 |

**Table S2.** Projections of the harvested rubber area increase required to meet the future demand projected by the International Rubber Study Group (IRSG) (IRSG 2022) to 2030 and reported by Tires & Parts News, January 19, 2022 (Tires & Parts News 2022). Assumes mean average yields between 2010 and 2020 from either Indonesia (0.89 t/ha) or Vietnam (1.69 t/ha) as reported in FAO statistics (FAO 2022). –Assuming yields of additional harvested area = 1.69 t/ha. ––Assuming yields of additional harvested area = 0.89 t/ha.

| Year | Observed natural rubber consumption (million metric tons) | Projected year-on-year consumption increase to following year (%) | Projected total natural rubber consumption (million metric tons) | Projected additional natural rubber consumption (million metric tons) | Min. additional area required *(million ha) | Max. additional area required **(million ha) |
| --- | --- | --- | --- | --- | --- | --- |
| 2020 | 12.7 | 9.4 | 12.7 | 0.0 | 0.0 | 0.0 |
| 2021 | 13.9 | 3.6 | 13.9 | 1.2 | 0.7 | 1.3 |
| 2022 | - | 2.3 | 14.4 | 1.7 | 1.0 | 1.9 |
| 2023 | - | 2.3 | 14.7 | 2.0 | 1.2 | 2.2 |
| 2024 | - | 2.3 | 15.0 | 2.4 | 1.4 | 2.6 |
| 2025 | - | 2.3 | 15.4 | 2.7 | 1.6 | 3.0 |
| 2026 | - | 2.3 | 15.7 | 3.1 | 1.8 | 3.4 |
| 2027 | - | 2.3 | 16.1 | 3.4 | 2.0 | 3.8 |
| 2028 | - | 2.3 | 16.5 | 3.8 | 2.2 | 4.2 |
| 2029 | - | 2.3 | 16.9 | 4.2 | 2.5 | 4.6 |
| 2030 | - | - | 17.2 | 4.6 | 2.7 | 5.1 |

**Table S3.** Projections of rubber production by 2030 using a simple linear model of production in response to year using FAOSTAT data for 2000 – 2020 (p < 0.0001, F = 805, d.f. = 19, adjusted r-squared = 0.98, beta = 407,046, meaning each year production increased by a predicted 407,046 metric tons). Production is then converted to required additional plantation area using mean average yields between 2010 and 2020 from either Indonesia (0.89 t/ha) or Vietnam (1.69 t/ha) as reported in FAO statistics (FAO 2022). –Assuming yields of additional harvested area = 1.69 t/ha. ––Assuming yields of additional harvested area = 0.89 t/ha. Code available in Supplementary Code 1, and input data in Data S4.

| Observed production (million metric tons) | Year | Fitted value for production (million metric tons) | Lower 95% CI | Upper 95% CI | Projected additional natural rubber consumption (million t) | Min. additional area required *(million ha) | Max. additional area required **(million ha) |
| --- | --- | --- | --- | --- | --- | --- | --- |
| 7.09 | 2000 | 7.3 | 7.0 | 7.6 |  |  |  |
| 7.48 | 2001 | 7.7 | 7.4 | 8.0 |  |  |  |
| 7.72 | 2002 | 8.1 | 7.9 | 8.4 |  |  |  |
| 8.41 | 2003 | 8.5 | 8.3 | 8.8 |  |  |  |
| 9.26 | 2004 | 8.9 | 8.7 | 9.2 |  |  |  |
| 9.52 | 2005 | 9.4 | 9.1 | 9.6 |  |  |  |
| 10.4 | 2006 | 9.8 | 9.6 | 10.0 |  |  |  |
| 10.6 | 2007 | 10.2 | 10.0 | 10.4 |  |  |  |
| 10.8 | 2008 | 10.6 | 10.4 | 10.8 |  |  |  |
| 10.3 | 2009 | 11.0 | 10.8 | 11.2 |  |  |  |
| 10.8 | 2010 | 11.4 | 11.2 | 11.6 |  |  |  |
| 11.6 | 2011 | 11.8 | 11.6 | 12.0 |  |  |  |
| 12.7 | 2012 | 12.2 | 12.0 | 12.4 |  |  |  |
| 13.1 | 2013 | 12.6 | 12.4 | 12.8 |  |  |  |
| 13.4 | 2014 | 13.0 | 12.8 | 13.2 |  |  |  |
| 13.3 | 2015 | 13.4 | 13.2 | 13.6 |  |  |  |
| 13.6 | 2016 | 13.8 | 13.6 | 14.1 |  |  |  |
| 14.4 | 2017 | 14.2 | 14.0 | 14.5 |  |  |  |
| 14.8 | 2018 | 14.6 | 14.4 | 14.9 |  |  |  |
| 15.0 | 2019 | 15.1 | 14.7 | 15.4 |  |  |  |
| 14.8 | 2020 | 15.5 | 15.1 | 15.8 |  |  |  |
| Predicted | 2021 | 15.9 | 15.5 | 16.2 | 1.0 | 0.6 | 1.2 |
| Predicted | 2022 | 16.3 | 15.9 | 16.6 | 1.4 | 0.8 | 1.6 |
| Predicted | 2023 | 16.7 | 16.3 | 17.1 | 1.8 | 1.1 | 2.1 |
| Predicted | 2024 | 17.1 | 16.7 | 17.5 | 2.2 | 1.3 | 2.3 |
| Predicted | 2025 | 17.5 | 17.0 | 17.9 | 2.7 | 1.6 | 3.0 |
| Predicted | 2026 | 17.9 | 17.4 | 18.4 | 3.1 | 1.8 | 3.4 |
| Predicted | 2027 | 18.3 | 17.8 | 18.8 | 3.5 | 2.1 | 3.9 |
| Predicted | 2028 | 18.7 | 18.2 | 19.2 | 3.9 | 2.3 | 4.4 |
| Predicted | 2029 | 19.1 | 18.6 | 19.7 | 4.3 | 2.5 | 4.8 |
| Predicted | 2030 | 19.5 | 18.9 | 20.1 | 4.7 | 2.8 | 5.3 |

## Supplementary Code 1

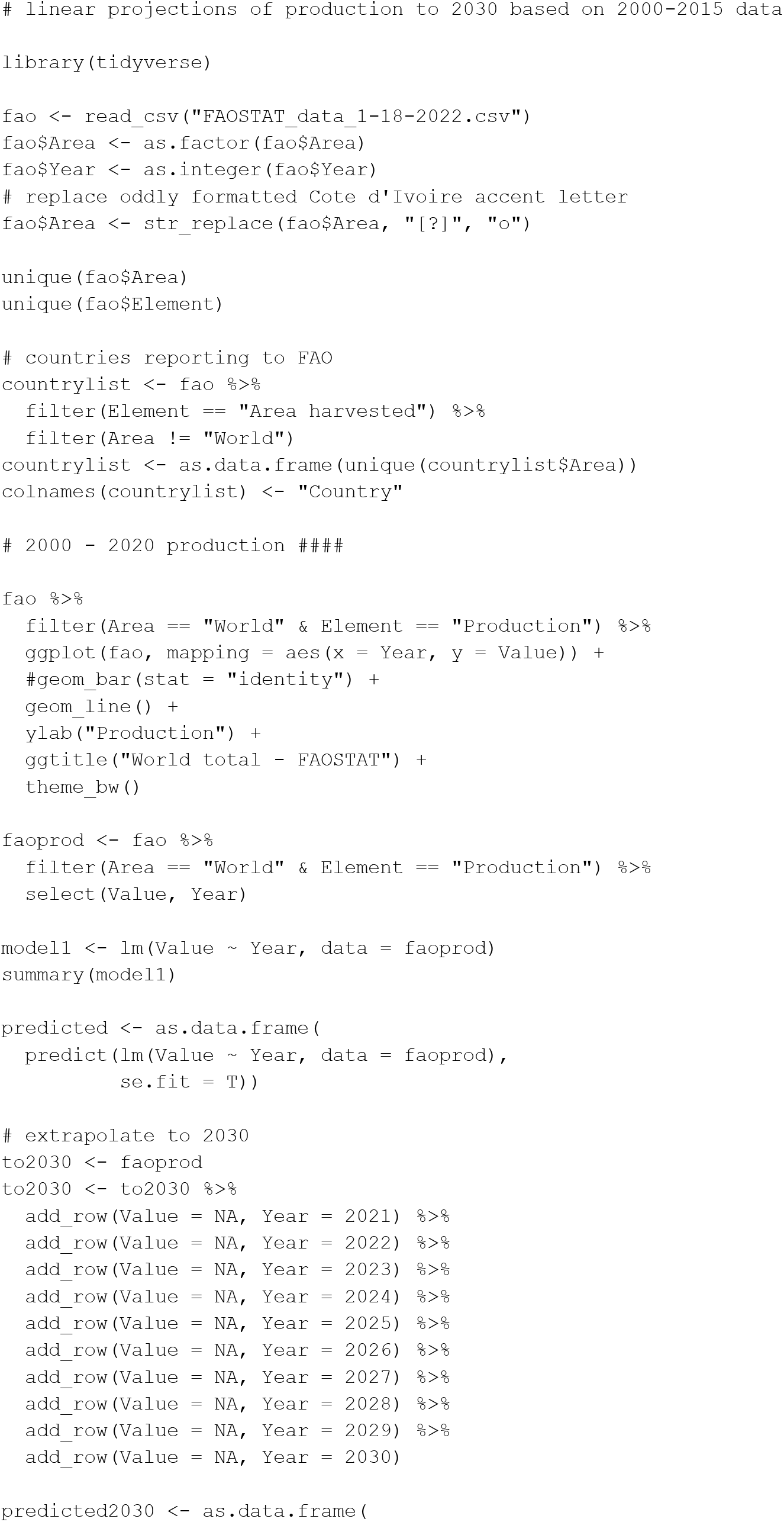
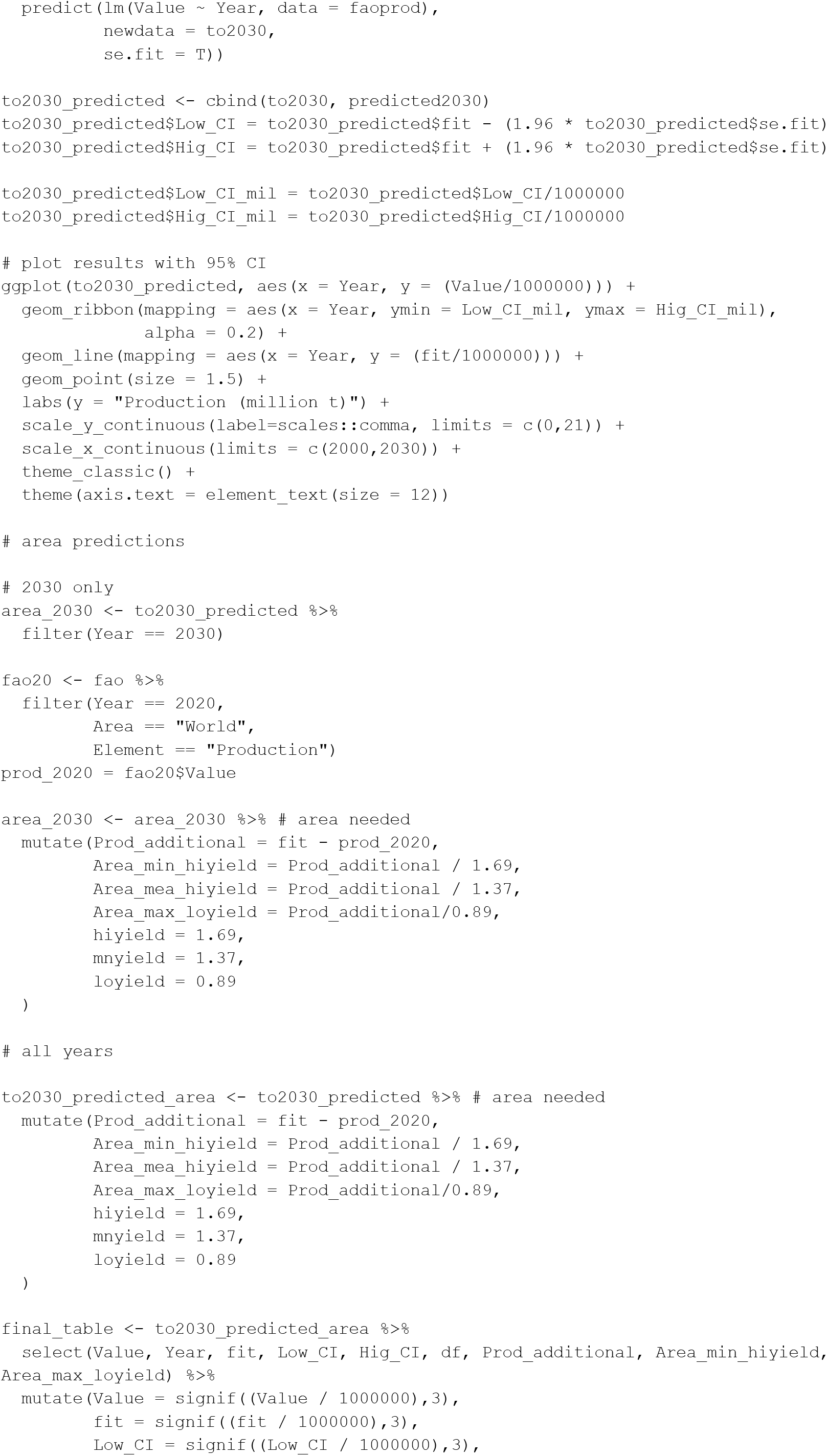
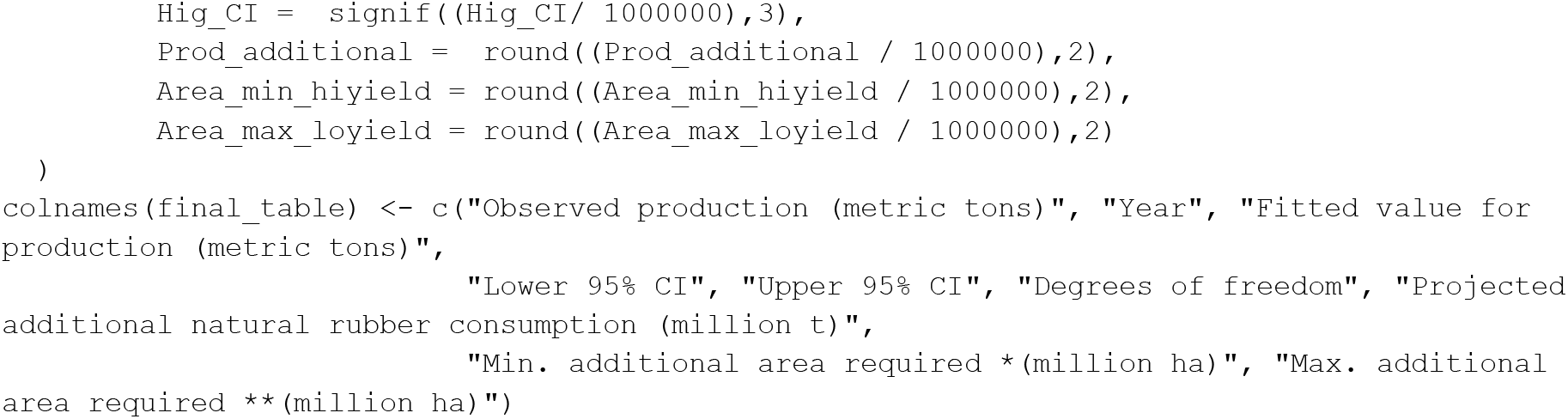

